# IL-33 Involved in the Progression of Liver Fibrosis Through Innate Immunity Cells (Mφ) Regulated by ICOS/ICOSL Signaling in Early Stage of Mice Schistosomiasis

**DOI:** 10.1101/2022.09.05.506595

**Authors:** Lei Liu, Peng Wang, Shi-Qi Xie, Wen-Jie Pu, Jing Xu, Chao-Ming Xia

## Abstract

Schistosomiasis liver fibrosis is characterized by the foci of a classical type 2 granulomatous inflammation in response to egg soluble antigen. The inducible costimulator (ICOS)/ICOSL signaling can mediate Th2 polarization and regulate the disease process of chronic schistosomiasis by adaptive immune response. ICOS/ICOSL signaling could regulate innate immune cells, including B cells and macrophage (Mφ), and participate in pulmonary fibrosis. The purpose of the study was to identify whether ICOS/ICOSL signaling could regulate the polarization of Mφ, and to explore the potential role of IL-33 in ICOSL knockout (ICOSL-KO) mice with schistosomiasis. Firstly, the expression level of CD86, CD206 and IL-33 was checked respectively in ICOSL-KO and WT mice infected with *Schistosoma japonicum* (*S. japonicum*). Then recombinant IL-33 (rIL-33) was injected into ICOSL-KO mice infected with *S. japonicum*. Our experiments showed that the liver Mφ was successfully polarized toward to classical activated Mφ (M1) and the expression of IL-33 was inhibited in ICOSL-KO mice. Furthermore, the injection of rIL-33 successfully aggravated liver fibrosis in ICOSL-KO mice, increased the numbers of lymphocyte antigen 6C (Ly6C)^hi^, enhanced the expression of C-C chemokine ligand (CCL)2,, CCL5 and C-X-C motif chemokine 2 (CXCL2), and promoted polarization of T helper (Th) cells to Th2 cells, as well as induced the autophagy and apoptosis of hepatic stellate cells (HSCs). Overall, the liver fibrosis was alleviated in ICOSL-KO mice along with the reduced expression of IL-33, which could skew the polarization of Mφ toward to M1, induce Th cells activation, HSCs apoptosis and autophagy through Smad2/3 and TGF-β signaling pathway, thus participated in the homeostasis of liver fibrosis of schistosomiasis.

## Introduction

Liver fibrosis is a chronic liver injury caused by a variety of factors, including viral infection, schistosomiasis, alcoholism [1]. To wound healing, liver fibrosis is an irreversible process. It is a key cause of liver cirrhosis in the early stage, and advanced liver fibrosis may eventually develop into liver cancer [2, 3]. Up to date, there are no specific and effective options in clinical treatment of liver fibrosis, it is urgent to discover novel disease-specific targets for antifibrotic purpose [3, 4]. Schistosomiasis is a zoonosis caused by *schistosoma* infection [5]. The main pathological mechanism of schistosomiasis is that the body immune system protects tissues and cells from the continuing and strong immune response caused by soluble egg antigen (SEA) of *S. japonicum*, through forming granuloma and fibrosis [6]. During the progression of schistosomiasis, a variety of inflammatory cells, such as macrophage (Mφ) and neutrophil, were activated and recruited around the granuloma. They stimulate hepatic stellate cells (HSCs) to proliferate to fibroblasts with the deposition of a large amount of extracellular matrix (ECM) [7]. When exceeding the metabolic capacity of the tissue, they would form the secondary fibrosis and cirrhosis [6].

Interleukin-33 (IL-33), as a novel member of IL-1 superfamily, is a kind of pro-inflammatory cytokine. It expresses in fibroblast-like cells, Mφ, endothelial cells and epithelial cells, and plays roles in regulation of homeostasis and inflammation [8]. The suppression of tumorigenicity (ST)-2, also known as IL-1 receptor ligand-1 (IL-1RL1, T1 or IL33R), is mainly expressed in fibroblasts, group 2 innate lymphoid cells (ILC2s), eosinophils and T helper 2 (Th2) cells [9]. Through binding to its receptor ST2, IL-33 shows crucial role in regulating type-2 immune response. It was reported that IL-33/ST2 has the ability to regulate the polarization of macrophage toward M2, subsequently producing type 2 chemokines, further actively inducing M2 to recruit inflammatory cells from micro-environment to regulate the immune response [10].

Inducible costimulator (ICOS) is identified as the third member of the CD28 family, expressing on activated T cells and resting memory T cell [11]. ICOS ligand (ICOSL), as the ligand of ICOS, also known as B7-related protein-1 (B7RP-1), LICOS, and GL50, is weakly expressed on the antigen-presenting cells at the steady state. Once these cells activated, the expression of ICOSL was observably upregulated [12]. ICOSL expressed not only on T cells of adaptive immunity, but also on Mφ and dendritic cells (DCs), which contribute to innate immunity [13, 14]. Many studies have reported that ICOS/ICOSL signaling pathway plays an important role in the adaptive immune response. Additionally, our previous study indicated that ICOS/ICOSL signaling regulates Th1/Th2 immune response, affecting Tfh differentiation in liver fibrosis process of schistosomiasis [15, 16]. However, the effects of ICOS/ICOSL on Mφ polarization and its precise mechanism in schistosomiasis remain unclarified. In this study, the expression of IL-33 decreased in ICOSL-KO mice. We hypothesized that IL-33 may play an important role in liver fibrosis of schistosomiasis, which rely on the ICOS/ICOSL singaling. We built the model of ICOSL-KO mice, infected them with *S. japonicum*, and injected recombinant IL-33 (rIL-33) to them. We found that rIL-33 successfully aggravated liver fibrosis in ICOSL-KO mice. Furthermore, rIL-33 injection presented higher numbers of Ly6C^hi^, enhanced the number of chemokines, promoted the polarization of Th cell to Th2, and finally affected the autophagy and apoptosis of HSCs. Moreover, we firstly found that Mφ in ICOSL-KO mice polarized to M1-type, thus reduced the severity of liver fibrosis. Therefore, our findings suggest that IL-33 participate in the liver fibrosis of mice schistosomiasis through regulating Mφ in innate immunity by ICOS/ICOSL signaling pathway.

## Results

### Pathological progression of schistosomiasis was attenuated in ICOSL-KO mice

Liver tissues were obtained for biochemistry test and pathological evaluation. Our data revealed that ICOSL-KO mice developed less severe phenotype than WT mice did 4 weeks post infection, such as less symptom and nodules. The weight of the liver tissue was decreased 8 weeks post infection in ICOSL-KO mice compared with that of WT mice (Supplementary Fig 1a, b). Histological examination was done to examine the area of granulomas and to evaluate the content of fibrosis. Compared with WT mice, the size of granulomas was significantly reduced in ICOSL-KO mice at 4, 8 and 12 weeks of post-infection (all *P*<0.05) (Supplementary Fig 1c, d). Furthermore, ICOSL-KO mice showed attenuate content of liver fibrosis as compared with WT mice at 8 and 12 weeks of post-infection (all *P*<0.05) (Supplementary Fig 1e, f). Moreover, we found that the content of liver HYP and the level of serum hyaluronic acid (HA) were decreased in ICOSL-KO group in comparison with WT group (*P*<0.05) (Supplementary Fig 1g, h). Since HYP and HA are important markers for evaluating the degree of liver fibrosis, these findings suggested that the block of ICOS/ICOSL signaling could effectively ameliorate liver fibrosis in mice infected with *S. japonicum*.

### ICOSL-KO mice upregulated the expression of M1

ICOSL is mainly expressed in B cells and Mφ, and the polarization of Mφ is related to liver fibrosis. Given the prior evidence, we hypothesized that ICOS/ICOSL signaling could also regulate the polarization of Mφ, which is involved in the progression of liver fibrosis. To test our hypothesis, we isolated the liver Mφ from ICOSL-KO/WT groups by density-gradient centrifugation to identify the expression of M1 and M2. As expected, the numbers of M1 and M2 showed no difference between WT and ICOSL-KO mice before infection. However, the numbers of M1 and M2 increased rapidly from week 4 till week 8 post infection and decreased at 12 weeks post-infection. M1 increased from week 4 to week 8 after infection in ICOSL-KO mice compared with that of WT mice. But M2 decreased from week 4 to week 8 after infection in ICOSL-KO mice compared with that of WT mice (Fig 1a-c). RT-PCR analysis showed that the expression of iNOS increased, while that of Arg1 decreased in ICOSL-KO mice, *P*<0.05 compared to that of WT mice (Fig 1d, e).

**Fig. 1.**
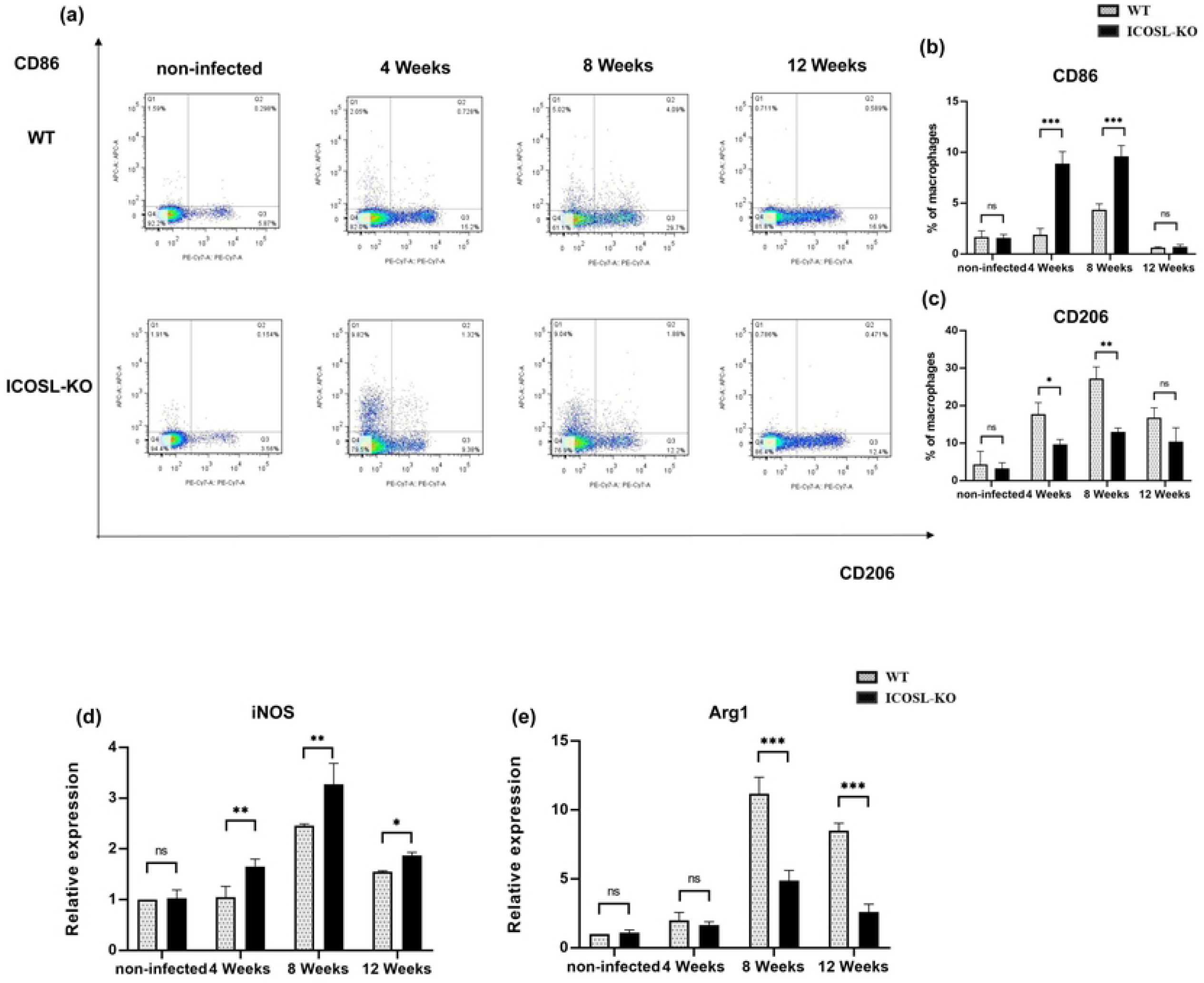
Lack of ICOSL increased the expression of CD86 in macrophages. (a-c) The liver macrophages were collected from *S. japonicum* infected mice by density gradient centrifugation and purified according to surface marker of F4/80 and CD11b by flow cytometry (FCM). The expression of CD86 and CD206 in the liver macrophage were determined by FCM (from two experiments with 4 mice per group). (d, e) The expression of *iNOS* and *Arg1* in the liver macrophage were determined by qRT-PCR (from two experiments with 4 mice per group). Data are presented as the mean ± SD from multi-group experiments. Statistical significance was calculated using independent t-test. \**p* < 0.05, \*\**p* < 0.01, and \*\*\**p* < 0.001

### Expression of IL-33 reduced in ICOSL-KO mice

IL-33 is a pro-inflammatory cytokine, playing crucial roles in regulating type-2 immune response [10]. These immune responses are associated with liver fibrosis in schistosomiasis [17, 18]. Along with shrunken size of granulomas and content of fibrosis in the liver of ICOSL-KO mice, the expression of IL-33 and ST2 significantly declined in murine schistosomiasis (Supplementary Fig 2a-f). The above data proved that ICOSL-KO mice have protective effect against liver fibrosis by inhibiting the expression of IL-33.

### IL-33 aggravated the liver fibrosis in ICOSL-KO mice

To investigate if IL-33 take part in the progression of *S. japonicum* hepatic pathology in ICOSL-KO mice, we injected exogenous IL-33 or recombinant IL-33, 1 μg/injection, once every week, totally five times, to ICOSL-KO mice after *S. japonicum* infection [19, 20]. Samples were harvested at the indicated times (Fig 2a). We found that the size of the hepatic granulomas and the degree of liver fibrosis were greater in rIL-33 injected mice than in control mice (*P*<0.05) (Fig 2b-e). Similarly, rIL-33 not only enhanced collagen deposition, but also increased hydroxyproline levels in 6 weeks, 8 weeks and 10 weeks after infection compared with those in control mice (*P*<0.05) (Fig 2f). This manifested that rIL-33 injection aggravated hepatic fibrosis in ICOSL-KO mice with *S. japonicum*. In addition, HA levels in serum were significantly higher in rIL-33 injected mice than that of control mice (*P*<0.05) (Fig 2g). Taken together, these data revealed an important role of IL-33 in aggravating *S. japonicum* egg-induced granuloma, hepatic injury, and hepatic fibrosis in ICOSL-KO mice.

**Fig. 2.**
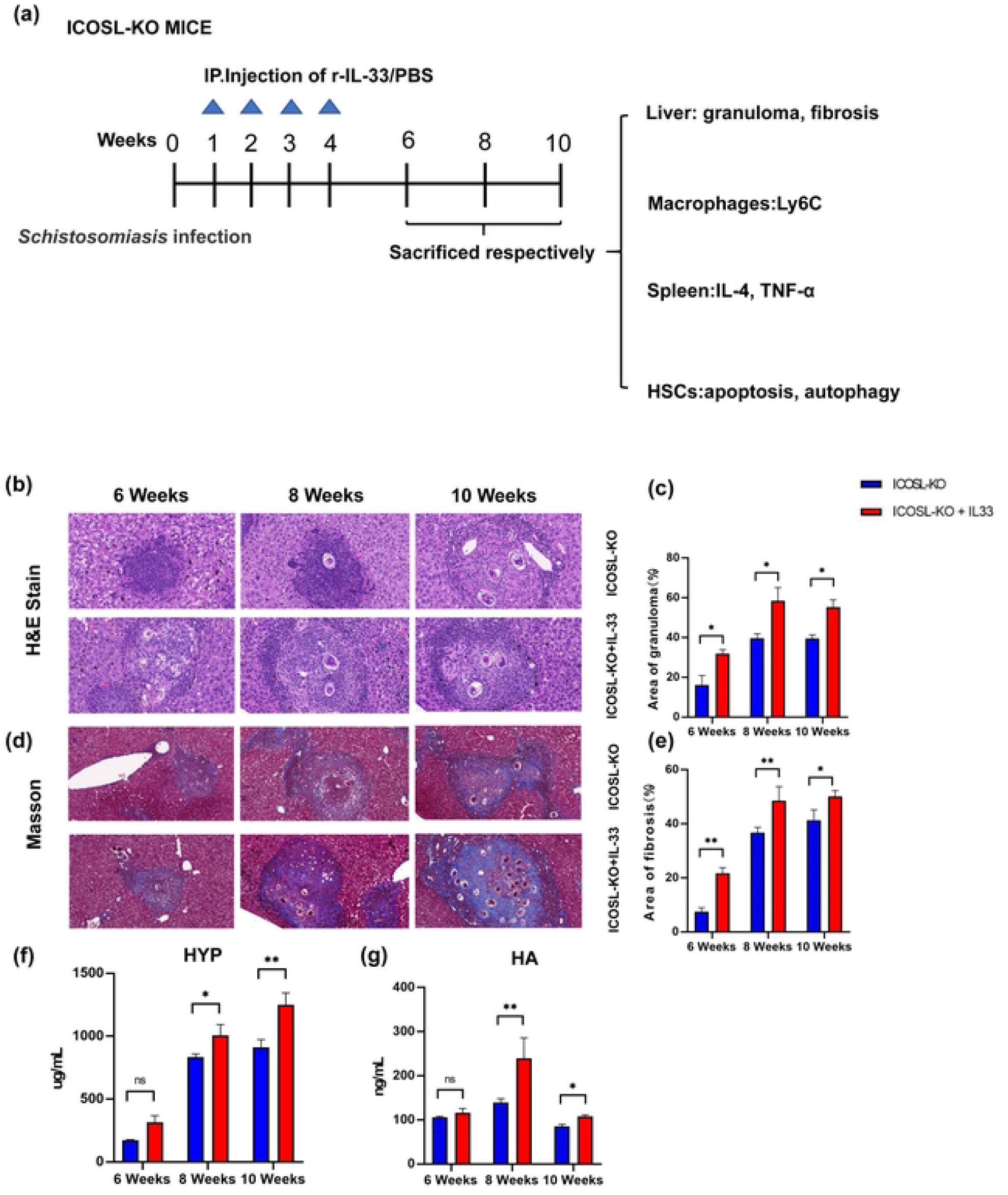
ICOSL-KO mice injected with rIL-33 exhibit more severe liver pathology than WT mice after *S. japonicum* infection. (a) Study design. rIL-33 mice and control mice were infected with 14±1 cercariae of *S. japonicum*, and liver samples from these mice were harvested at the times indicated after infection (from two experiments with 25 mice per group). Representative graphs of H&E staining (b, c) and Masson staining (d, e) of liver specimens (from two experiments with 4 mice per group), scale bars, 200 μm. (f) The HYP content in the liver was measured by hydroxyproline assay kit (from two experiments with 4 mice per group). (g) Sera hyaluronic acid level was tested by enzyme-linked immunosorbent assay (from two experiments with 4 mice per group). Data are presented as the mean ± SD from multi-group experiments. Statistical significance was calculated using independent t-test. \**p* < 0.05, and \*\**p* < 0.01

### The expression of F4/80 and Ly6C increased in the liver of rIL-33 injection mice after *S. japonicum* infection

To assess whether IL-33 participate in the suppressive effect of restorative Ly6C^lo^ monocytes after *S. japonicum* infection, we checked the number of circulating monocytes to estimate the levels of chemokines that promote Mφ recruitment, and detected the number of restorative Ly6C^lo^ monocytes and proinflammatory Ly6C^hi^ monocytes in ICOSL-KO mice injected with rIL-33 at the acute stage of schistosomiasis infection. As expected, IL-33 treated mice harbored more Ly6C^hi^ and enhanced the chemokine levels. As illustrated in Fig 3a, b, the total number of circulating monocytes increased in rIL-33 group than that in control group. The results of flow cytometric showed that Kupffer cells (F4/80^hi^, CD11b^inter^) increased in rIL-33 group compared with that of control group, whereas the number of infiltrating Mφ (F4/80^inter^, CD11b^hi^) decreased (*P*<0.05) (Fig 3c-e). Simultaneously, infiltrating monocytes were further identified as two distinct subsets: proinflammatory Ly6C^hi^ monocytes and restorative Ly6C^lo^ monocytes. Through flow cytometric analyses, we found that more proinflammatory Ly6C^hi^ monocytes penetrated into the liver of rIL-33 mice than of control mice after infection (*P*<0.05). By contrast, the numbers of Ly6C^lo^ monocytes decreased (*P*<0.05). We also analyzed the chemokines production in rIL-33 mice by qRT-PCR and found that administration the expression of CCL2, CCL5 and CXCL2 significantly increased after rIL-33 injection (Figure 3f-h). These results indicated that IL-33 partake in the fibrosis activity by recruiting pro-inflammatory Ly6C^hi^ monocytes and secreting CCL2, CCL5 and CXCL2.

**Fig. 3.**
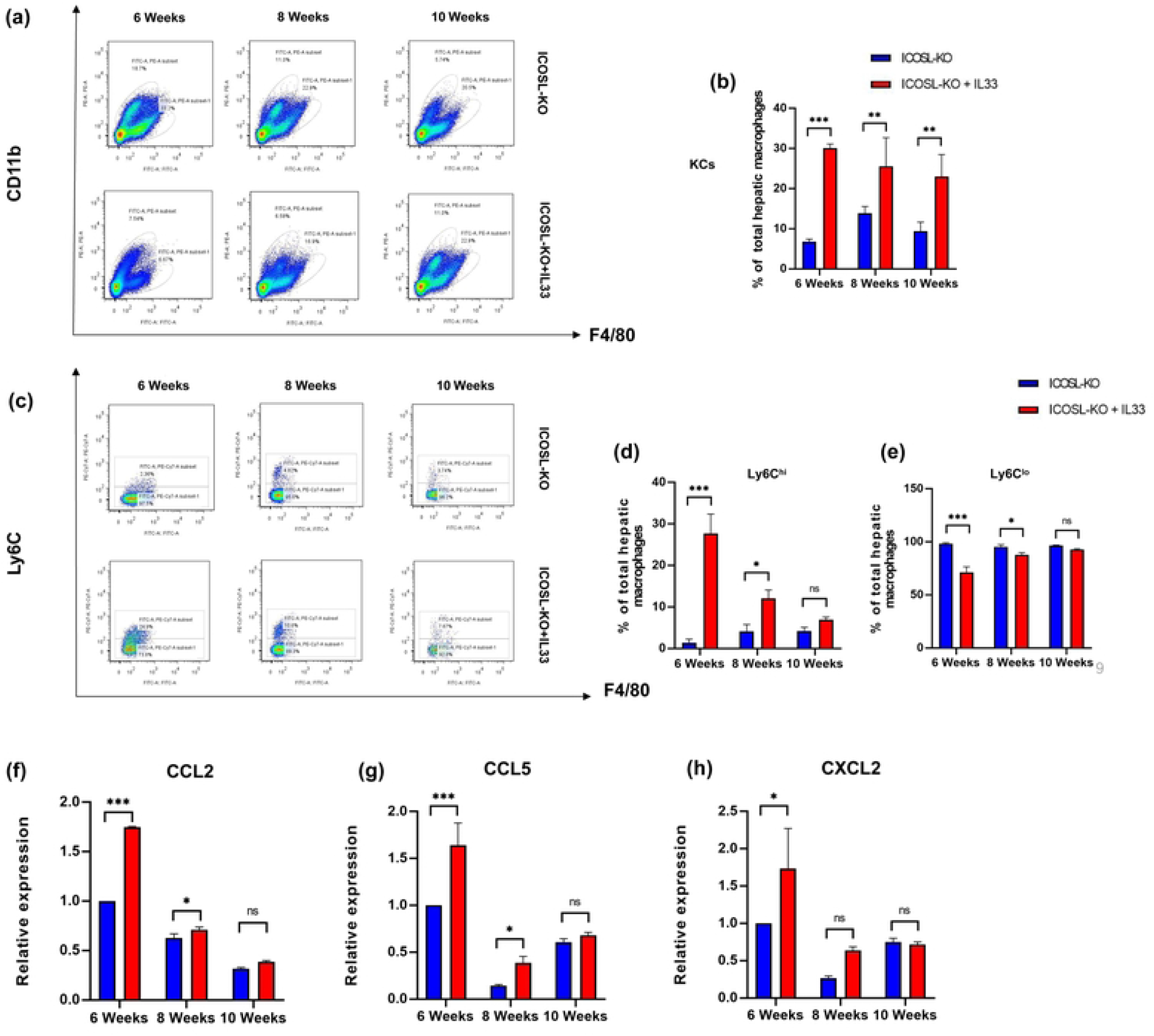
rIL-33 injection causes increase of macrophages in the liver. (a, b) The infiltration of hepatic macrophages (CD11b+F4/80+) after injection with rIL-33 were quantified by flow cytometric analysis (from two experiments with 4 mice per group). (c-e) Graphical summary displayed the percentages of KCs (CD11b^lo^F4/80^hi^), Ly6C^hi^ monocytes (CD11b^hi^F4/80^lo^ly6C^hi^), and Ly6Clo monocytes (CD11b^hi^F4/80^lo^ly6C^lo^) out of total hepatic macrophages (CD11b+F4/80+). Relative numbers of KCs, Ly6C^hi^ monocytes, and Ly6C^lo^ monocytes in rIL-33 injection mice and control mice (from two experiments with 4 mice per group). (f–h) The RNA of liver macrophages was extracted to analyze the mRNA expression of several chemokines, including *CCL2, CXCL2* and *CCL5* (from two experiments with 4 mice per group). Data are presented as the mean± SD from multi-group experiments. Statistical significance was calculated using independent t-test. \**p* < 0.05, \*\**p* < 0.01

### rIL-33 participated in liver fibrosis partly by regulating Th1, Th2 in *S. japonicum* infected ICOSL-KO mice

Given the interaction of IL-33 with Th2 cells in adaptive immunity, we explored the effect of rIL-33 on the cytokine secretion of Th2 cells after *S. japonicum* infection. We isolated splenic CD4+T cells from rIL-33 treated mice and control mice, and assessed the changes of Th1 and Th2 in spleens by flow cytometry. We observed a dramatic reduction of the TNF-α percentage from week-6 after *S. japonicum* infection, while that of IL-4 increased from week-8 in rIL-33 treated mice compared with in control mice (*P*<0.05) (Figure 4a-d).

**Fig. 4.**
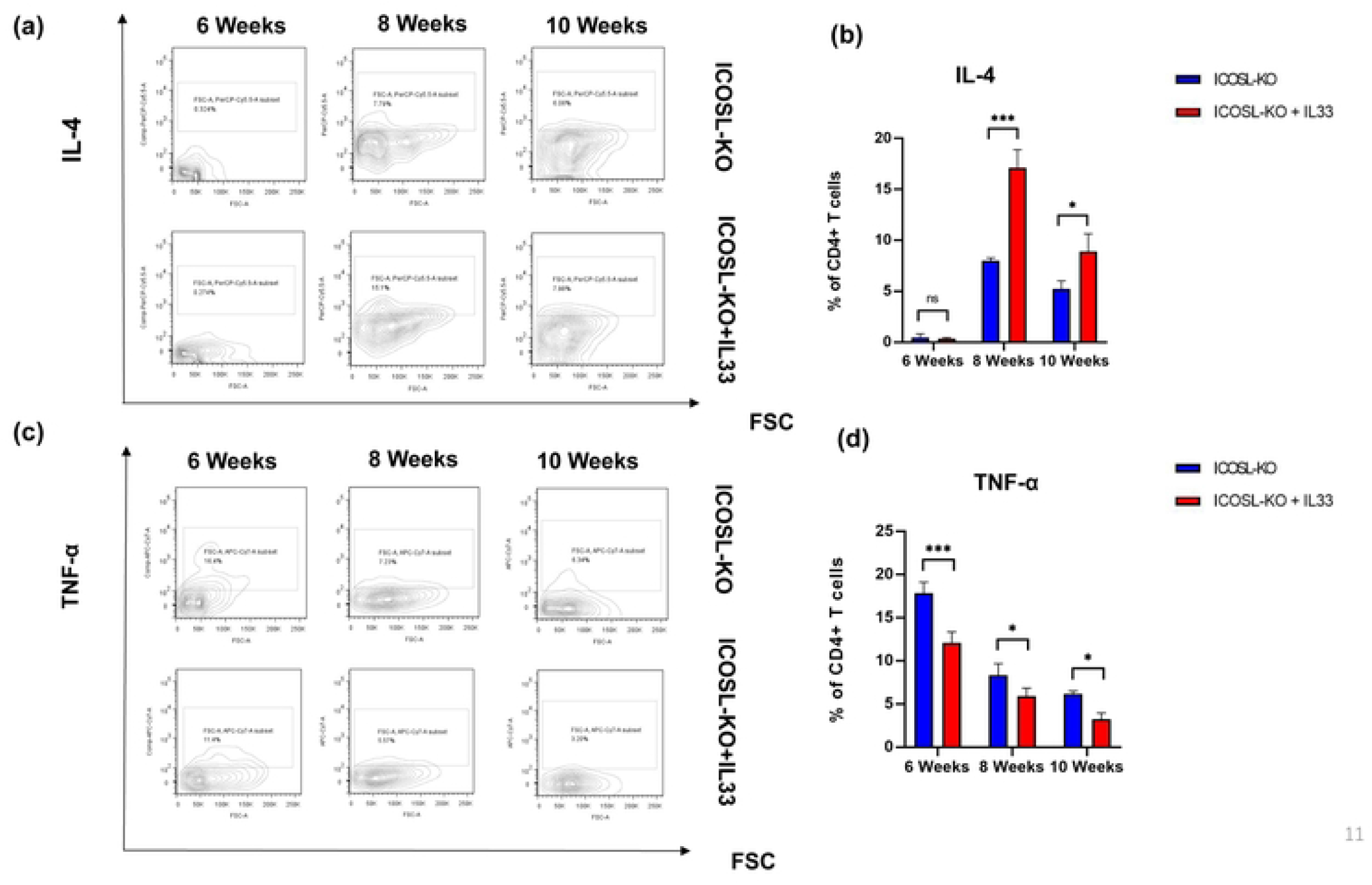
Lower expression of IL-4 on the surface of CD4+ T cells in ICOSL-KO mice infected with *S. japonicum*. The spleen cells were collected as described in the Materials and Methods, and then stained for CD4. The percentages of IL-4 (a, b) (from two experiments with 4 mice per group) and TNF-α (c, d) (from two experiments with 4 mice per group) are analyzed by FCM. Data are presented as the mean ± SD from multi-group experiments. Statistical significance was calculated using independent t-test. \**p* < 0.05, \*\**p* < 0.01

### rIL-33 promoted the apoptosis and autophagy of HSCs

To unearth the function of IL-33 on HSCs, we examined the apoptosis rate of JS1. Our data showed that the apoptosis rate of activated HSCs significantly decreased in rIL-33 group than in control group (Fig 5a, b). Additionally, we analyzed the role of IL-33 in the autophagy of HSCs. As expected, the autophagy rate of activated HSCs significantly increased in the rIL-33 treated group compared with in control group (Fig 5c, d).

**Fig. 5.**
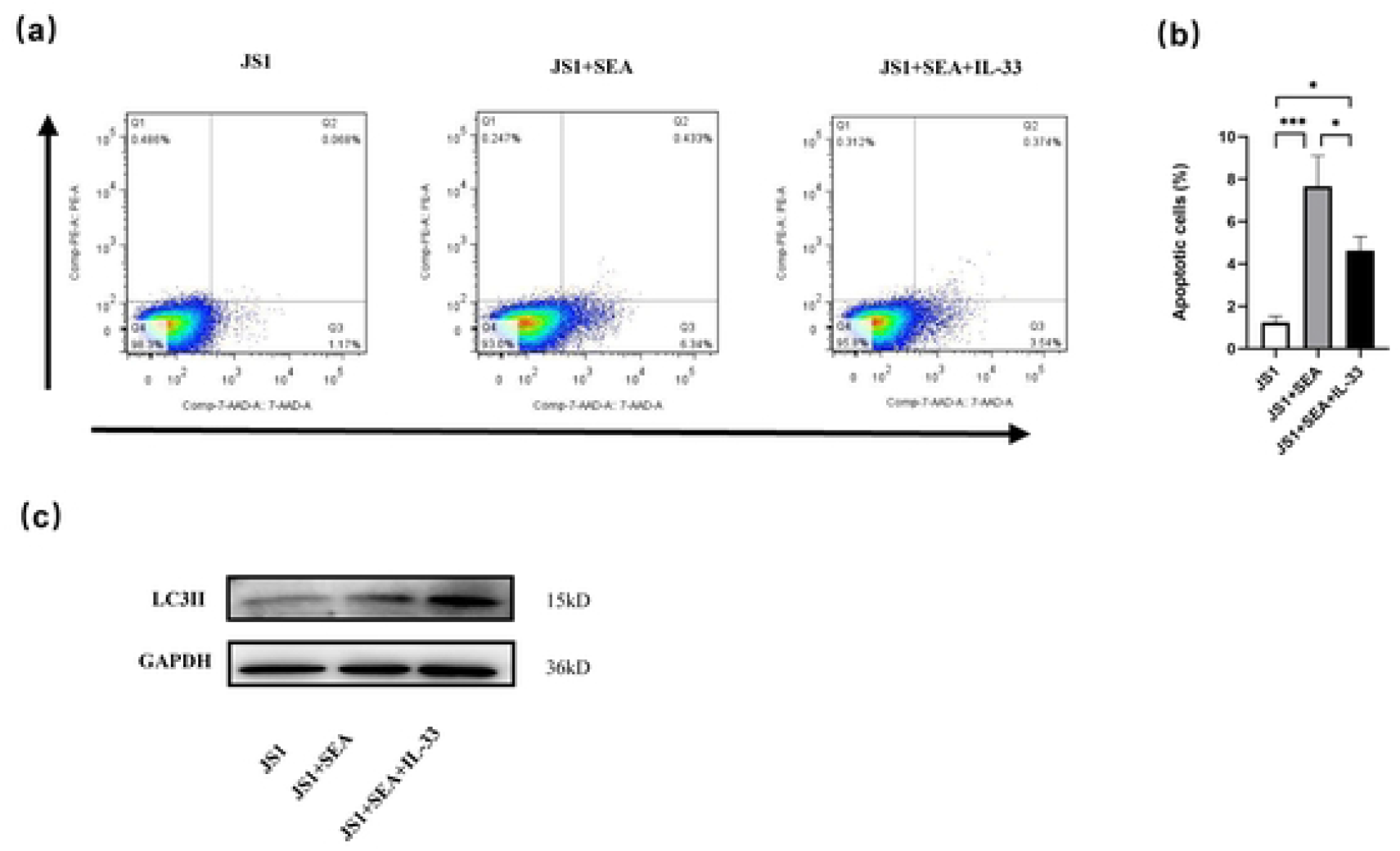
rIL-33 promotes the apoptosis and autophagy of HSCs. (a, b) HSCs (JS1) were treated with PBS or IL-33 (15 ng/mL) for 72 h, respectively, and then JS1 cells were collected and assessed for apoptosis by FCM (from three experiments per group). (c) The autophagy of JS1 was detected by WB (from three experiments per group). Data are presented as the mean ± SD from multi-group experiments. Statistical significance was calculated using independent t-test. \*\**p* < 0.01

### rIL-33 aggravated liver fibrosis via Smad2/3/TGF-β1 axis

IL-33 was positively correlated with TGF-β1 expression, and IL-13 and TGF-β1 expression was decreased in the IL-33 -/- mouse model [21]. In addition, TGF-β1/SMAD pathway has been identified as a major pathway that promote inflammatory response in schistosomiasis by activating HSCs [21]. To verify whether IL-33 exacerbated liver fibrosis is regulated by the Smad2/3/TGF-β1 axes in the infected mice, we compared the expression of Smad2/3/TGF-β1 in rIL-33 infected mice with in control mice. The results showed that the expression of TGF-β1 and Smad2/3 was both up-regulated in the schistosome-infected livers after rIL-33 treatment (Figure 6a-f). As reported, TGF-β1/Smad2/3 signaling pathway are strongly activated during liver fibrosis. TGF-β1-mediated phosphorylation of Smad2/3 not only contributes to the activation of HSCs, but also is correlated with the apoptosis of activated HSCs [22, 23]. Our findings above are well in accordance with this.

**Fig. 6.**
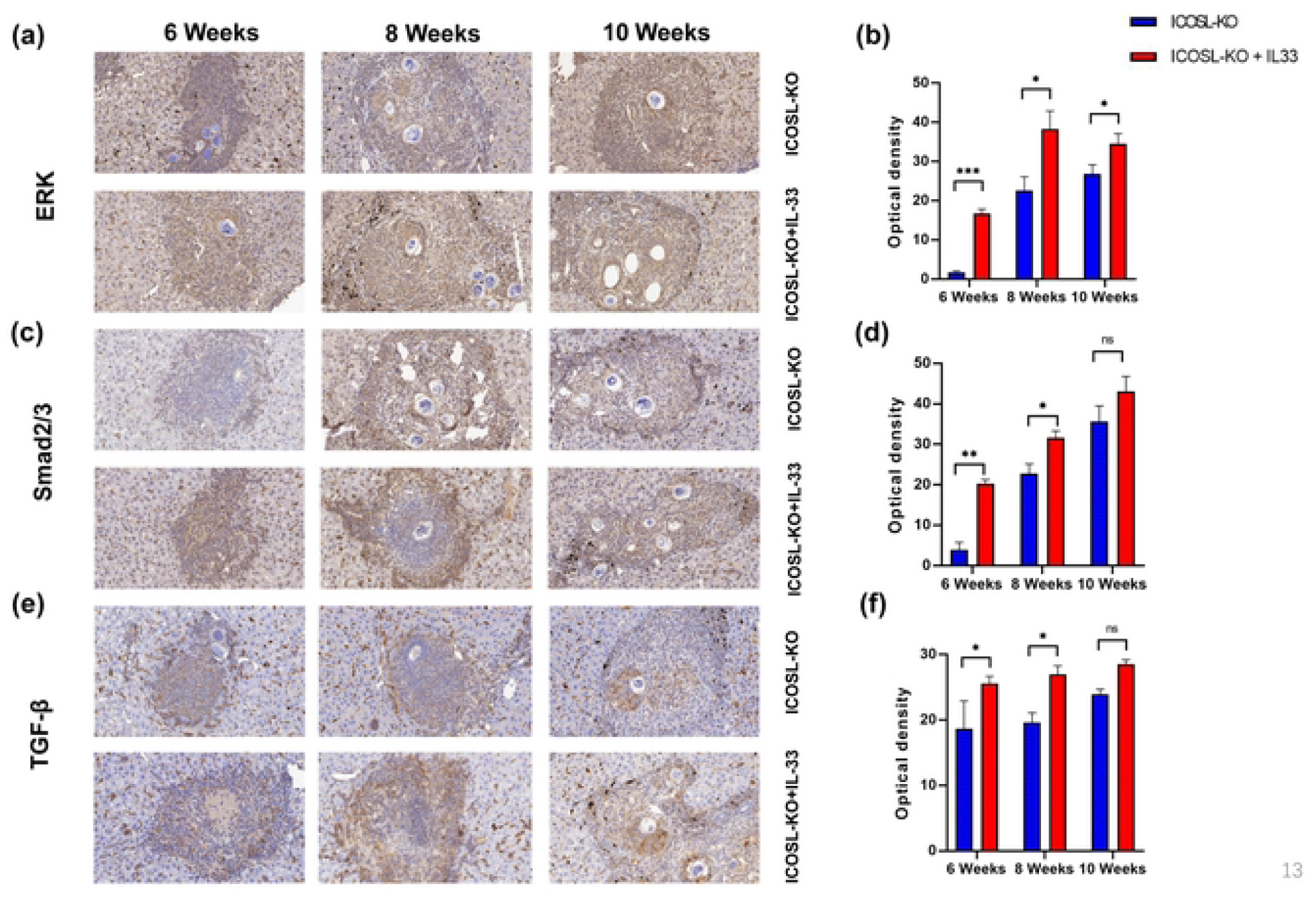
The expression of fibrosis-related factors in ICOSL-KO and WT mice. (a-f) To explore the potential mechanism of rIL-33 on the exacerbation of liver pathology in ICOSL-KO mice after *S. japonicum* infection, we measured the expression of ERK, SMAD2/3 and TGF-β by immunohistochemistry (from two experiments with 4 mice per group). Data are presented as the mean ± SD from multi-group experiments. Statistical significance was calculated using independent t-test. \*\**p* < 0.01

## Discussion

As the second signaling, ICOS/ICOSL costimulatory signaling is necessary for the homeostasis and function of various immune cell populations, and plays different roles in a series of disease models [24–26]. Studies have found that blocking ICOS/ICOSL costimulatory signaling could inhibit immunoreactions, reduce the symptoms of G6PI-induced arthritis, and lighten the severity of collagen-induced arthritis by inhibiting the Th effect response in *vivo* [27]. Whereas, some studies have suggested that blocking ICOS/ICOSL costimulatory signaling in Treg cells promoted the deterioration of diabetes symptoms [28]. Many scientific studies reported that the ICOS/ICOSL signaling pathway plays an important role in the adaptive immune response [13, 29]. However, ICOSL expresses on antigen-presenting cell, such as dendritic cells and Mφ in innate immunity, and their expression is closely related to fibrosis in many tissues [11]. Tanaka *et al*. suggested that ICOSL expression on B cells and Mφ was inversely associated with the severity of Bleomycin-Induced lung and skin fibrosis [30]. However, the effects of ICOSL on the polarization of Mφ in the hepatic fibrosis progression of schistosomiasis are still unclear.

To assess the role of ICOS/ICOSL interaction in schistosomiasis liver fibrosis, we compared the immune responses and pathological reactions in ICOSL-KO and WT mice infected with *S. japonicum*. In this study, we found that ICOSL-KO mice infected with *S. japonicum* showed fewer granuloma and less liver fibrosis than that in WT mice. Meantime, ICOSL-KO-Mφ presented a M1-dominated cell phenotype, which suggested that blocking of ICOS/ICOSL signaling is crucial in regulating the polarization of Mφ to M1. To our knowledge, this is the first report that ICOS/ICOSL signaling could affect the polarization of Mφ, and then alleviate the liver fibrosis of schistosomiasis. Therefore, our findings proved that ICOS/ICOSL signaling contributes to against schistosomiasis hepatic fibrosis through regulating innate immunity at the early stage of *S. japonicum* infection in mice. What is interesting, the expression of IL-33 decreased in ICOSL-KO mice, which suggested there is a relation between IL-33 and the liver fibrosis in ICOSL-KO mice.

Through binding to its receptor ST2, IL-33 shows crucial roles in regulating type-2 immune response [31]. The schistosomal liver fibrosis was also reported to associated with Th2 polarization, caused by Th2-cell-type cytokines in response to allergen exposures [32]. IL-33 not only amplifies Th2 cytokines (IL-4, IL-5 and IL-13), but also shifts the polarization of Mφ to M2, leading to higher infection intensity and liver immunopathology [33, 34]. During the pathological process of *S. japonicum* infection, the phenotype of Mφ shifted to M1-type inflammation at the early stage, but to M2-type at the acute stage. This restricted the secretion of soluble antigens by *schistosome* eggs, and meanwhile aggravated the content of granulomas and liver fibrosis during the process. Therefore, different phenotypes of Mφ play diverse roles in pathogens resisting, tissue repairing and recovering of liver fibrosis during the process of chronic schistosomiasis. Since liver fibrosis in schistosomiasis involves strong immune components, we applied this model to ICOSL-KO mice to investigate the roles of these molecules in modulating innate and adaptive immunity. IL-33 has been considered to be an important cytokine to adjust the immunity effect of Th2/M2, and it contributes to HSCs activation and collagen deposition in liver fibrosis [35]. We premised that the Mφ regulated by ICOS/ICOSL could adjust the expression of IL-33, and then affect the development of liver fibrosis. Studies have reported that ICOS/ICOSL signaling can affect IL-33 through PSTAT5 and Bcl-2, and its blocking can significantly decrease the levels of IL-13 and IL-5 [36]. However, how IL-33 affects the liver pathological process of schistosomiasis is still unclear. As IL-33 had been previously shown to induce macrophage-mediated *S. japonicum-induced* hepatic pathology mice [33], we applied the ICOSL-KO-based model to test the functional requirements for IL-33 in cytokine production and functions of macrophage. Plasma IL-33 level was suggested to negatively correlate with schistosomiasis infection [37]. We observed that IL-33 and ST2 expressed less in Mφ of ICOSL-KO mice than of WT mice. Accordingly, we speculated that there are some correlations between Mφ and IL-33 in ICOSL-KO model of schistosomiasis. Thus, we injected rIL-33 to ICOSL-KO mice infected with *S. japonicum* to evaluate the phenotype of Mφ and verify the effect of IL-33-predominant immune response on hepatic fibrosis. In rIL-33 treated group, the liver pathology became significantly aggravated, judged from the areas of fibrosis, HYP in the liver, and HA level in serum.These results demonstrated that IL-33-dominant immunity had significant impact on the hepatic granulomas and fibrosis.

It has been reported that resident Kupffer cells in murine liver are F4/80^hi^CD11b^intermediate^, whereas recruited monocyte-derived Mφ are CD11b^hi^F4/80^intermediate^ [38, 39]. In the current study, we found rIL-33 injection in ICOSL-KO mice increased the numbers of resident Kupffer cells and monocyte-derived Mφ. Ly6C has been widely used to identify functionally distinct populations of circulating murine monocytes and Mφ in diseased tissues: Ly6C^hi^ monocytes/macrophages are thought to initiate inflammatory activities, whereas Ly6C^lo^ monocytes/macrophages promote healing [40, 41]. In the present study, Mφ expressed high levels of Ly6C^hi^, indicating that they belong to a pro-inflammatory subset. ICOSL-KO-Mφ was believed to present a M1-dominated cell phenotype. Once we injected rIL-33 into ICOSL-KO mice, the Mφ shifted to monocyte-derived Mφ to Ly6C^hi^, which aggravate schistosomal liver fibrosis.

In addition, IL-33 also induces Mφ to release CCL2 that could recruit monocytes from blood into liver and then enhance ly6C^hi^ recruitment. Thus, IL-33 contributes to fibrosis formation by regulating matrix deposition. It is generally accepted that inflammation promotes fibrosis and liver disease progression [42, 43]. Interestingly, our data showed that more Mφ infiltrated in the liver of rIL-33 injected mice and liver fibrosis increased after *S. japonicum* infection. Macrophages have been shown to respond to lipid insults by producing pro-inflammatory mediators and matrix-degrading enzymes that putatively regulate liver fibrosis. These mediators include TGF-β, matrix metallopeptidases (MMP)2, MMP7, MMP9, MMP12 and MMP13, *etc*. Many chemokine genes, including CXCL2, CCL2, CCL3, CCL4, and CCL5, were upregulated in rIL-33 treated Mφ compared with in control Mφ. Macrophage-secreted chemokines can recruit innate and adaptive immune cells in the liver.

Macrophages and HSCs are the main cell types involved in the pathogenesis of hepatic granulomas [44]. IL-33 can not only acts indirectly on the process of liver fibrosis by affecting the polarization of Mφ, but also directly contributes to collagen deposition by promoting the activation of HSCs into myofibroblasts. IL-33 produces TGF-β1 that directly promotes HSCs differentiation into fibrogenic-SMA positive myofibroblasts (activated HSCs) through TGF-β1-Smad2/3 signaling [6, 45]. IL-33 indirectly activates NF-κB and MAPK that regulate the apoptosis and autophagy of HSCs. IL-33 regulates the activation of HSCs and produces copious amounts of IL-13, aggravating the liver fibrosis during infection [45]. *In vitro*, rIL-33 induces the activation of HSC through the p38 MAPK pathway, which is mediated by ERK, JNK and p38 protein kinase, and can activate and regulate the autophagy of HSCs. Moreover, as a soluble cytokine, IL-33 is produced by activated HSC and stimulates pro-fibrogenic Th2 cytokines which is suggested to be the leading producers of ECM proteins [42]. IL-33 bind to ST2, play important role in diverse inflammatory diseases. At the end-stage of infection, *S. japonicum-induced* liver fibrosis is associated with elevated ST2 levels [46]. IL-33/ST2 regulates the signaling of TLR4 and NF-κB to affect the expression of MMP9 and tissue inhibitors of metalloproteinases (TIMP)-1, and finally affect collagen deposition [47, 48]. Yet, our study indicated that rIL-33 injection lead to a high expression of TGF-β and Smad2/3, which are closely related to the polarization of Mφ [49, 50], the activation of Th cells [51–53], and the apoptosis and autophagy of HSCs [54, 55]. When compared with that of control group, rIL-33 promoted Mφ skew to Ly6C^hi^, mediated the activation from T cell to Th2, increased autophagy of HSCs and inhibited the apoptosis of HSCs. Thus, it is plausible that IL-33 secretes TGF-β and Smad2/3, both participating in the liver fibrosis of schistosomiasis.

In summary, our study firstly explored the cellular and molecular mechanisms of IL-33 mediated by ICOS/ICOSL in the deterioration of schistosomal liver fibrosis. We hypothesized that ICOS/ICOSL signaling as a check point in promoting the polarization of innate immunity (Mφ) to M1, which led to lower expression of IL-33 in limiting schistosomiasis liver fibrosis. We injected rIL-33 into the ICOSL-KO mice infected with *S. japonicum*. Our studies showed that the liver granulomas and fibrosis were significantly exacerbation after rIL-33 injection. Furthermore, rIL-33 injection greatly increased the secretion of TGF-β1 and Smad2/3, and promoted Mφ skew to Ly6C^hi^, stimulated T cell activating to Th2, increased autophagy of HSCs and inhibited the apoptosis of HSCs in experimental mouse liver fibrosis. Our investigation also suggested that Mφ response in the liver may rebalance *S. japonicum*-triggered immunity, minimize granulomas formation and alleviate fibrotic process in ICOSL-KO mice. Our findings open a possibility of IL-33 as a therapeutic candidate, through regulating the innate immunity (Mφ) by ICOS/ICOSL signaling to treat the schistosomal liver fibrosis (Supplementary Fig 3).

## Ethics approval

Animal experiments were carried out in strict accordance with the Regulations for the Administration of Affairs Concerning Experimental Animals (1988.11.1), and all efforts were made to minimize animal suffering. All animal procedures were approved by the Institutional Animal Care and Use Committee (IACUC) of Soochow University for the use of laboratory animals (Permit Number: 201604A136). The Institutional Ethical Committee of Soochow University and all experiments were conducted in accordance with the principles of the Declaration of Helsinki including any relevant details.

## Availability of data and material

The data and material that support the findings of this study are available from the corresponding author on reasonable request.

## Consent to participate

All the study subjects provided informed consent to participate in this study.

## Authors’ Contributions

L. L., P. W. and C. X. designed and conducted the experiments; L. L., P. W. and S. X; contributed to development of methodology; L. L., P. W., and W. P. contributed to analysis and interpretation of data; L. L., F. D., and J. X., prepared figures and wrote the manuscript; C. X. supervised the project and edited the manuscript.

## Materials and Methods

### Ethics statement

All mice experiments were conducted according to the Institutional Animal Care and Use Committee (IACUC) of Soochow University for the use of laboratory animals (Permit Number: 2007–13), and all efforts were made to minimize suffering.

### Mice, parasites, and infection

Fifty ICOSL-KO mice and fifty wild-type mice, six-to eight-week-old, female, were used for mouse experiments. ICOSL-KO mice were purchased from Jackson Laboratory (Bar Harbor, Maine, USA) and WT mice were purchased from Shanghai SLAC Laboratory animal co. LTD (Shanghai, China). All mice were raised under specific pathogen-free conditions at the laboratory animal research facility of Soochow University (Suzhou, China). Snails (*Oncomelania hupensis*) harboring *S. japonicum* cercariae (Chinese mainland strain) were purchased from Jiangsu Institute for Schistosomiasis Control (Wuxi, China). Each mouse was infected with 14±1 cercariae of *S, japonicum* through the abdominal skin. At 4, 8 and 12 weeks of post-infection, ten mice were randomly allocated as the experimental group respectively, while ten non-infected mice were kept as control group and sacrificed for further studies.

### Histology assay

The left upper lobe of each liver was harvested from experimental groups and control groups, fixed with 10% formalin for 48 h, and then embedded in paraffin following standard procedure. About 4-μm-thick paraffin sections were dewaxed and incubated according to the manufacturer’s instructions. Embedded liver tissues were stained with hematoxylin and eosin (H&E) or MASSON and photographed under bright-field images. The size of hepatic egg granulomas (five per mouse) was measured, reflecting hepatic fibrosis degree. All images were analyzed with computer image analysis system (Image-Pro Plus software, Media Cybernetics, Inc., Rockville, MD, USA).

### Hydroxyproline (HYP)

HYP content in the liver was measured according to the instructions of the hydroxyproline assay kit. The absorbance was measured at 550 nm on an ELISA reader.

### Quantitative-Real time PCR (qRT-PCR)

Total RNA of frozen liver tissues or cells were extracted using Trizol (Invitrogen, Carlsbad, CA, USA), according to the manufacturer’s instructions. The first strand cDNA was synthesized using a High-capacity cDNA Reverse Transcription kit (Invitrogen, Carlsbad, CA). qRT-PCR was performed by a QuantStudio 6 Flex real-time PCR detection system (Applied Biosystems, Foster City, CA) with SYBR Premix Ex TaqTM II (Tli RNaseH Plus) (TaKaRa, Beijing, China) and Universal PCR Master Mix (Applied Biosystems™) according to the manufacturer’s instructions. The primers were shown in Table 1. After qRT-PCR, threshold cycle (C_T_) values were obtained and then applied to the 2^−ΔΔ C_t_^ method for the calculation of gene relative expression level. All experiments were repeated three times.

### Western Blotting

Samples were homogenized in Radio Immunoprecipitation Assay buffer containing a cocktail of protease inhibitors (Santa Cruz, CA). Protein extracts were loaded onto 8/15% polyacrylamide gels and then transferred onto a PVDF membrane (Millipore, Burlington, MA). Membrane was then blocked by incubating in 5% skim milk/TBS for 1 h at RT, and incubated with primary antibodies (Abs) and HRP-conjugated secondary Abs (Bioss, Beijing, China). Protein bands were visualized with ECL-chemiluminescent kit (Millipore, MA). The Abs against LC3B was purchased from Novusbio (Littleton, CO). The Abs against GAPDH were purchased from Abcam (Cambridge, MA).

### Liver Macrophages isolation

Liver macrophages were isolated by perfusing liver with 50%/25% Percoll (GE Healthcare, Pittsburgh, PA). Single-cell suspension of macrophages were incubated with Fc-blocker (2.4 G2; BD PharMingen) and Zombie Aqua dye (Biolegend, CA) for 10 minutes. Macrophages were then stained with antibodies of interest for 30 minutes at 4°C in the dark. The following antibodies were used: anti-CD11b (Biogems, CA), anti-F4/80 (Biolegend, CA), anti-CD206 (BD, CA), anti-CD86 (Biosciences, CA), anti-Ly6C (BD, CA). Flow cytometry analysis was performed using the FCM Calibur (BD FACSVerse system, CA).

### HSCs isolation

HSCs were isolated by perfusing liver with 15%/11.5% Optiprep (Axis-shield), final cell concentration was around 1×10^6^ cells/100 μl PBS. The apoptosis of cells was stained with PE Annexin V Apoptosis Detection Kit (BD Biosciences, CA, USA) according to the manufacturer’s instructions. To stimulate cells, HSCs were cultured and treated with bovine serum albumin (BSA) (10%), BSA (10%)/IL-33 (15 ng/ml), Palmitic acid (PA) (0.1 mM), or Palmitic acid (0.1 mM).

### Splenocyte culture

Single-cell splenocyte suspensions were prepared by mincing the spleens in PBS (Sigma Corporation, St. Louis) containing 1% FBS (Gibco, Grand Island, NY). Red blood cells were lysed using ACK lysis buffer. Soluble egg antigens (SEA) of *S. japonicum* were purchased from Jiangsu Institute for Schistosomiasis Control (Wuxi, China), with a concentration of 25 μg/ml of SEA. The splenocytes from infected mice were cultured in RPMI 1640 medium (Gibco, Grand Island, NY) containing 10% FBS, 100 U of penicillin/ml, and 0.1 mg/ml of streptomycin. For IL-4 and TNF-α staining, the cells were firstly cultured for 4 h in a 37°C/5% CO_2_ incubator with 50 ng/ml PMA (Sigma) and 1 mM ionomycin (Sigma, Burbank), and then surface-stained with fluorescein isothiocyanate-conjugated anti-mouse CD4 antibody (Millipore, Billerica) for 30 min at 4°C in the dark. Subsequently, the cells were fixed and permeabilized with Cytofix/Cytoperm buffer (Millipore, Billerica), and then intracellularly stained with Peridinin chlorophyll protein-Cy5.5-conjugated anti-mouse TNF-α (Millipore, Billerica) and phycoerythrin-conjugated anti-mouse IL-4 (eBioscience, San Diego, CA).

## Acknowledgments

We thank the study participants as well as the staff involved in the collection of blood samples. We appreciate Dr. Fuhong Dai at Medical College of Soochow University to provide the English language editing.

---

Supplementary Fig. 1 The alleviation of pathological progression of schistosomiasis in ICOSL-KO mice. (a, b) The mice were infected with 14±1 cercariae of *S. japonicum*. A total of 25 mice (five mice per group) were randomly allocated and sacrificed at 0 (before infection), 4, 8 or 12-weeks post-infection.

The appearance and weight of liver were exhibited (from two experiments with 4 mice per group). (c, d) Liver granuloma was identified by HE staining, and the granulomas area was measured with quantitative analysis (from two experiments with 4 mice per group). (e, f) Liver fibrosis as determined by Masson staining and liver fibrosis content was showed with quantitative analysis (from two experiments with 4 mice per group), scale bars, 200 μm. (g) Sera hyaluronic acid level was assayed by enzyme-linked immunosorbent assay (from two experiments with 4 mice per group). (h) Liver HYP was measured by the alkaline lysis method (from two experiments with 4 mice per group). Data are presented as the mean ± SD from multi-group experiments. Statistical significance was calculated using independent t-test. \**p* < 0.05, \*\**p* < 0.01, and \*\*\**p* < 0.001

Supplementary Fig. 2 The decreased expression of IL-33 in ICOSL-KO mice.

(a-f) Validation of the levels of IL-33 and ST2 in the liver of ICOSL-KO mice. The levels of IL-33 and ST2 in the liver were analyzed by immunohistochemistry (IHC), scale bars, 200 μm. The mRNA expression of IL-33 and ST2 were measured by qRT-PCR (from two experiments with 4 mice per group). Data are presented as the mean ± SD from multi-group experiments. Statistical significance was calculated using t-test. \**p* < 0.05, \*\**p* < 0.01, and \*\*\**p* < 0.001

Supplementary Fig. 3 The graphical abstract of cellular and molecular mechanisms of rIL-33 on the aggravation of severe liver pathology in ICOSL-KO mice of schistosomiasis.

